# Frisky CALF sometimes outruns LASSO

**DOI:** 10.1101/2020.03.27.011700

**Authors:** Clark D. Jeffries, John R. Ford, Jeffrey L. Tilson, Diana O. Perkins, Darius M. Bost, Dayne L. Filer, Kirk C. Wilhelmsen

## Abstract

Regularized regression analysis is a mature analytic approach to identify weighted sums of variables that predict outcomes. Typically, the number of subjects (N) is smaller than the number of predictors (p). Here, we present a novel coarse approximation linear function (CALF) to frugally select important predictors, build linear models, and discover causality.

CALF is a linear regression strategy applied to normalized data that employs only a few (such as 2 to 20) nonzero weights, each +1 or -1. Metrics can be Welch t-test p-value, area under curve (AUC) of receiver operating characteristic, or Pearson correlation, depending upon data type and user preferences. For quantitative approximations, a linear fit (adding an intercept value and rescaling ±1 weights by a common multiplier) can be added to optimize mean squared error (MSE).

Real medical data of five types were used to generate examples with goals of binary classification or real variable approximation. Predictors considered were real, sets of integers, or ternary values of single nucleotide polymorphisms. When applied to real data, CALF approximations outperformed in p-value, AUC, correlation, or MSE a popular regularized linear regression algorithm, namely, basic LASSO. It appears that using LASSO without considering CALF might risk wasting resources.

**Availability:** R version: Comprehensive R Archive (CRAN): https://cran.r-project.org/web/packages/CALF/index.html

Python 3.x version: GitHub: https://github.com/jorufo/CALF_Python

## Main

Regularized regression modeling strategies for choosing variables and weights are mature and extensively documented^1^. Ordinary (linear) least-squares (OLS) minimizes the sum of squared differences between the N real entries of a targeted dependent vector *vs*. a weighted linear combination of p predictors. Finding optimal weights is an application of algebra and calculus dating to the early 1800s and works of Legendre and Gauss. However, OLS can provide uninterpretable and unstable weights when exact or approximate collinearity exists between predictors or when problems are underdetermined (i.e., N << p). Regularized regression may employ Lagrange multipliers to constrain the optimization; examples include Tikhonov regularization (ridge regression, using constrained L2-norm), least absolute shrinkage and selection operator (LASSO, using constrained L1-norm), and elastic net (using constrained convex sum of the L1- and L2-norms)^1-4^. Each method has its advantages and disadvantages. As one advantage, basic LASSO with L1-norm includes choice of a parameter called s that can force a solution with few nonzero weights, enabling cross-referencing of selected predictors and a search for underlying meaning among them^5^. Thus, discovering small sets of collectively informative predictors may inform epistasis and suggest causal networks.

However, Hand argued that devising novel, exotic methods to solve a specific data classification or approximation problem until one finally outperforms another is a mere “illusion of progress”^6^. Therefore, establishing the utility of any novel regression model requires overcoming a substantial burden of proof. We understand that challenge and so provide diverse examples in which CALF succeeds as well as a mathematical analysis for why it should do so.

We present coarse approximation linear function (CALF), an algorithm that, applied to real data, builds models with frugal use of predictors, superior goodness of fit metric values *vs*. those of a gold standard, as well as superior permutation test performance. These properties could yield relatively simple causal interpretation of the interplay of predictors. Furthermore, the improvements in some cases could mean the difference between a discovery of statistical significance or a mere trend.

CALF assumes predictor values have been a priori cleaned and corrected for confounders and then converted to z-scores. Binary dependent vectors (e.g., control *vs*. case) are represented as 0 or 1 values, but other types of dependent vectors are converted to z-scores. For goodness of fit in terms of Welch t-test p-value (p-value), AUC, or Pearson correlation (correlation), this preprocessing enables a “coarse” selection of weight values, that is, a restriction of nonzero weight values to ±1. We show how this seemingly primitive approach can outperform basic LASSO. If MSE fit is desired for head-to-head LASSO comparisons, then a simple OLS calculation yields a linear conversion (adjustment) of the CALF solution, that is, an intercept value and a common multiplier to scale ±1 weights. This process does not alter p-value, AUC, or correlation, but optimizes MSE. Considering the widely acknowledged value of basic LASSO, we focus our comparisons with LASSO as the “gold standard.”

By using coarse weights, CALF may provide greater model stability in the sense that slight changes in input data (e.g., deletion of one subject) may change the ±1 weight estimates little or not at all. Also, if two or more predictors have very similar or identical values, CALF will simply employ at most one of them in a solution and ignore the rest. Generally, using CALF avoids collinearity issues.

CALF has already been applied in three longitudinal psychiatric studies to find sums of predictors measured at initial clinical presentation that collectively predict whether a newly presenting patient is likely to transition to psychosis within two years^7-9^. In those papers CALF was sketched and heavily used, but no underlying details were provided. As shown *infra*, an improved program now yields updates of those early findings.

## Methods

CALF uses a greedy forward selection algorithm to accumulate a limited number of nonzero weights, each ±1. A dependent vector Y may be compared to the product of an N-by-p data matrix X and a weight vector β with -1, 0, or +1 components in terms of a chosen metric (p-value, AUC, or correlation). In the initial step all β entries are 0, and one by one each is changed in the very first step to +1 (or possibly -1 to make correlation values positive). The metric values of the products as tentative solutions are noted and the first such selection that is at least as good as any subsequent selection is kept. In subsequent iterations the remaining 0 entries in β are changed one by one to ±1, and the first such selection, if any, that improves the goodness of fit and does so at least as well as any subsequent change is kept. Else, if at any iteration there is no such choice to improve the metric, then CALF ends. Else, if the number of nonzero β entries reaches a preset limit L, then CALF ends. Else, if no unselected predictors remain, then CALF ends. This is not the same as simply choosing predictors with low p-values. Also, as a greedy algorithm, CALF can become “stuck” on a local optimum and ignore a global optimum (Supplement 1).

In conventional regression methods, the comparison of dependent vector Y *vs*. Xβ is scored by MSE, but in basic CALF the goal is different. We seek metric values meaning: a Welch t-test p-value decreasing from default = 1; an AUC increasing from default = 0.5; or a correlation increasing from default = 0. Note that only the direction in p-space of the vector β matters for these three metrics; β can be rescaled by any positive multiplier without affecting them. The fact that only direction matters is an underlying reason that CALF works with simple ±1 weights. Thus, basic CALF is a very simple greedy algorithm. Pseudocode for CALF follows.

### Data

N-by-1 response vector Y (column matrix);

N-by-p predictor matrix X

### Parameters

natural number L ≤ p, a limit on the number of predictors to be employed in a solution.

## Result

p-by-1 weight vector β (column matrix) with at most L nonzero entries (weights) = ±1;

Let d^k^ denote the kth column of `diag`(1);

~~~
**begin**
       β ← 0;
       s ←0;
       for i ≤ min(L,p) **do**
                β’ ←0;
                s’← 0;
                **foreach** β_j_ ← 0 **do**
                       s^+^ ← score(βX + d^j^ X);
                       s^-^← score(β X + d^j^ X);
                       s’_j_ ← max(s^+^, s^-^);
‘
                       β’_j_ **if** s’_j_ = s^+^ **then** 1 **else** -1;
                **end**
                s_max_ max(s’);
                **if** s_max_ – s > Δs **then**
                        s← s_max_;
                        β ← 0;
                **end**
                **else break**
      **end**
**end**
~~~

If in addition a low MSE value for direct comparison with a LASSO solution is desired, then a basic CALF solution may be adjusted by OLS from CALF to adjCALF = b+m(Xβ) where m (a common, positive multiplier) and b (a single intercept value repeated in a column vector) are derived as scalars from OLS applied to Y and Xβ. That is, b is a N-by-1 matrix with all entries b, and m > 0 is real multiplier of all entries in Xβ.

For the sake of fairness, the LASSO solution (using MSE as metric) should also be adjusted. Adjusting the LASSO solution by OLS improves the LASSO MSE, preserves the number of nonzero weights in the LASSO solution, does nothing to its p-value, AUC, or correlation score, but violates the original L1-LASSO constraint on weights.

## Model selection and comparison

When evaluating models, we seek simplicity, goodness of fit, and permutation performance:

1. Simplicity: Select a set of at most 20 collectively informative predictors from a large set of potential predictors.
2. Goodness of fit: Seek good p-value, AUC, correlation, or MSE metric value, as appropriate or preferred.
3. Permutation performance: Compute an empirical p-value from permutation tests and reject algorithms above 0.05 as ineffective.

Also desirable is consistent marker popularity. That is, we may select many (e.g., 1000) random subsets of true data (e.g., 90%) and apply CALF to each with the limiting number of nonzero weights = 20. A histogram of selection counts may then reveal those predictors that are frequently selected (e.g., > 500 times). Avoiding infrequently selected predictors may be a way of avoiding overfitting and using predictors of small effect size. Ideally, the popularities of selected predictors should display a “cliff,” that is, a sharp decrease in popularities for predictors not in a small pool. Thus, a cliff might suggest an optimal L value, enabling the first goal.

The second goal assumes appreciation of the strengths and sometimes subtle shortcomings of the metrics. A full discussion of them is beyond the scope of the present paper, but goodness of fit of any metric in itself does not preclude overfitting.

The third goal—and also the first—may be served by examination of metric values in permutation tests. The empirical p-value (not to be confused with the metric p-value) herein is a type of probability value well-known in permutation testing^10^. The empirical p-value of a given algorithm relative to a given classification or approximation problem is an estimate of an upper limit of the risk that any apparent success (goodness of fit) has been achieved by chance.

Empirical p-value testing entails applying a given algorithm to many (e.g., 1000) pseudo-data versions of a classification or approximation problem in which all components of the dependent vector have been randomly permuted. Empirical p-value is computed from the empirical number E of times the algorithm performance is by chance superior in D versions of pseudo-data to performance of the algorithm using real data^11,12^; specifically, empirical p-value = (E+1)/(D+1).

Sweeping through L = 1, 2, …, 20 generally reveals a particular choice of L for CALF with minimal empirical p-value. If two choices have about the same empirical p-value, then the one with smaller L is preferred. Popularity histograms can also contribute to the selection of L. Similarly, sweeping through a range of s values in basic LASSO may reveal indirectly an optimal number of employed predictors for that algorithm.

If a value for the limit L in CALF is selected, then the particular CALF solution with L as limit is called CALFL. Likewise, LASSOs is defined. CALFL and LASSOs mean versions of the algorithms with L in the range 1, 2, …, 20 and s values that indirectly yield about the same range of numbers of nonzero weights.

### Five examples of data sources and their analyses

We will illustrate the performances of CALF and LASSO with five data sets drawn from three previously reported studies and two publicly available sources (a convenient site for all five is in out Implementation). Our goal is realistic mathematical illustration only—not generation of medical hypotheses requiring extensive knowledge of the diseases mentioned. For this reason, predictors not chosen in the models (GitHub address of data matrices in Implementation) are encoded simply as M1, M2,… parameters of data sets and algorithm performances are in Table 1.

**Table 1.** Parameters of five examples with and CALF and LASSO metric values.

| Example | # Subjects |  | # Predictors | Dependent | Predictor | CALF metric | LASSO metric |
| --- | --- | --- | --- | --- | --- | --- | --- |
|  | Controls | Cases |  | Vector |  |  |  |
| 1 | 40 | 32 | 135 | binary | Blood analyte levels | p-value | MSE |
| 2 | 37 | 30 | 130 | binary | Leukocytic miRNAs levels | AUC | MSE |
| 3 | 39 | 32 | 19 | binary | symptom severity | p-value | MSE |
| 4 | 213 | 165 | 140 | binary | Blood analyte levels | p-value | MSE |
| 5 | NA | 582 | 9312 | continuous | SNP allele coding | correlation | MSE |

Simple data cleaning and normalization methods were used on all our examples, and medical research would demand careful reconsideration of this step. In examples 1 and 2, averages and standard deviations of data for a set of unaffected persons were used to generate z-scores for each predictor. In the other examples (unaffected persons not included) all measurements of a given predictor were used to generate z-scores. The ages of onset data in the fifth example were converted to z-scores as well. Notably, in the fifth example the relatively large dimension of the dependent vector (582) allowed cross-validation testing, as shown *infra*.

## Results

### Example 1

We first utilized a dataset from the North American Psychosis-Risk Longitudinal Study^13^. Here the goal is use of blood plasma analyte levels to predict future development of psychosis in subjects meeting research criteria for psychosis high-risk. Data included levels of 135 blood analytes from 72 subjects, 40 who did not and 32 who did subsequently develop frank psychosis. The 135 blood analyte levels were determined from samples taken at the time of study enrollment^7^. The CALF solution here differs from that of the original publication^7^ which used a different metric.

The metric herein was p-value, so the very first predictor chosen by CALF was that with the lowest of 135 p-values. However, subsequent choices were not generally associated with rank of p-value among remaining predictors, an important aspect of collective classification. That is, initially selected predictors provide a sum that can be improved by additional predictors with nuanced discrimination power. The empirical p-values for L = 1 to 20 for Example 1 are shown in Figure 1(a). Another important CALF parameter is the popularity profile of predictors. We measured popularity by applying CALF20 to 1000 random 90% subsets of nonconverters and 90% subsets of converters. High popularity reflects a predictor with consistent importance (Figure 1(b)). Regarding LASSO solutions, 1(c) shows that as s sweeps from 0.155 (selecting one predictor) to 0.055 (selecting 19 predictors (no s value yields 20 predictors)), the best LASSO empirical p-value is 0.126.

**Figure 1.**
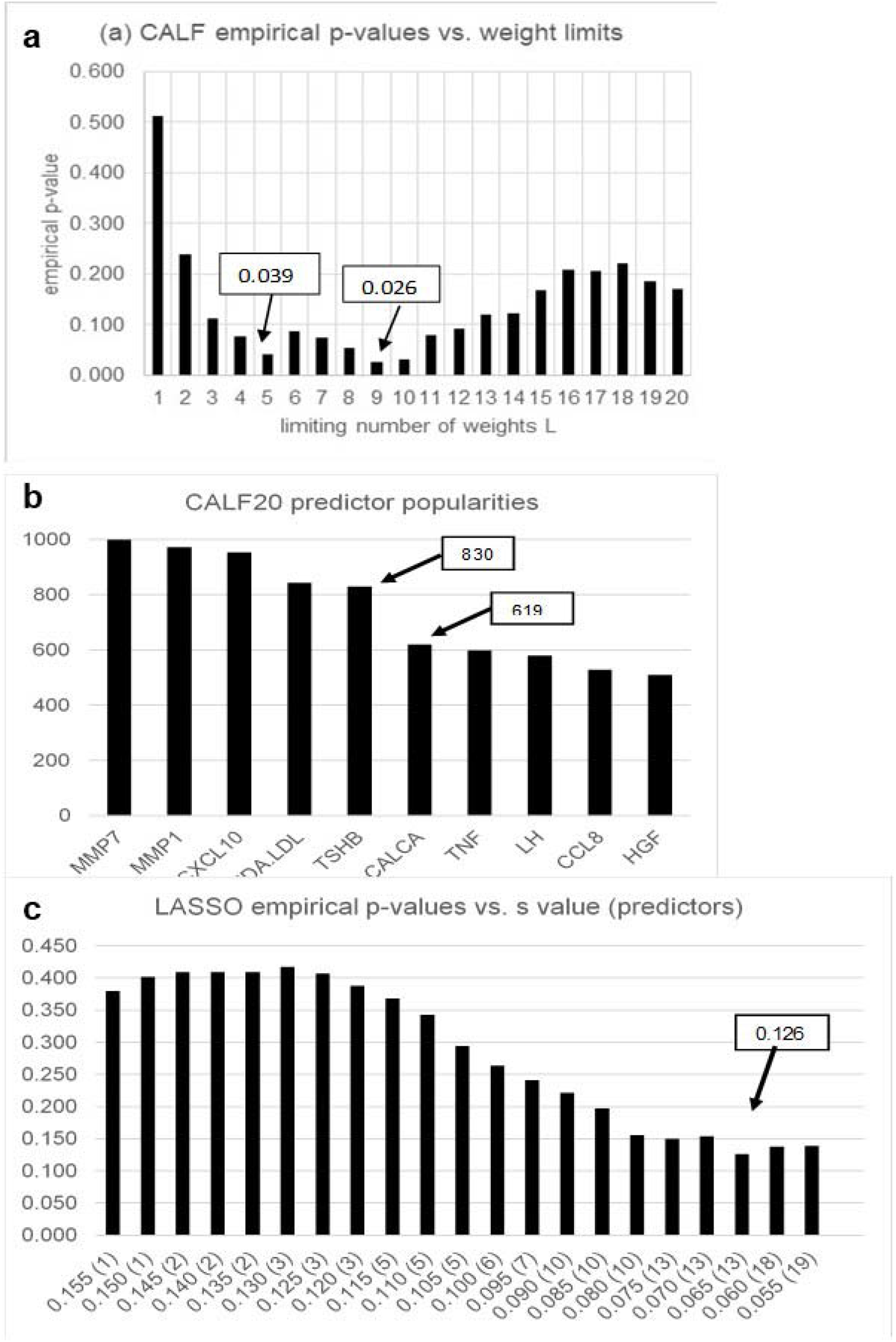
(a) For L = 5, empirical p-value (arrow) is 0.039 (Welch p-value = 2.55E-9); empirical p-value of CALF9 is smaller (0.026). However, (b) shows a cliff at five predictors in the popularity profile of the top 10 selected by CALF20. Thus, simplicity and the cliff strongly recommend CALF5. Meanwhile, (c) shows that as s sweeps from 0.155 (selecting one predictor) to 0.055 (selecting 19 predictors (no s value yields 20 predictors), LASSOs achieves a lowest empirical p-value of 0.126 (at s = 0.065, selecting 13 predictors).

In more detail, we consider the solutions from CALF and LASSO

CALF5 = +MMP7 +MDA-LDL - MMP1 +TSHB - CXCL1

LASSO.065 = +0.398 +0.121*MMP7 -0.059*MMP1 +0.059*MDA.LDL +0.050*TSHB -0.048*CXCL10 +0.040*FTL+0.021*CCL8+0.017I*GHE+0.013*APOD+0.007*KITLG0.006*IL6 +0.003*TTR +0.003*IL1B

(The analyte symbols are defined in the earlier publication^7^. There are no s values for which LASSOs selects 4, 8, 9, 11, 12 14, 15, 16, 17, or 20 predictors. The real weights in the shown LASSO.065 are approximated to three only decimal places.)

### Comparison

The empirical p-values (0.039 *vs*. 0.126) and the optimized number of predictors (5 *vs*. 13) for CALF5 and LASSO.065 imply CALF is a superior algorithm, given the constraints that at most 20 predictors are used and the metric is p-value.

In Supplement 2 the same example was examined by the following eight classifier construction methods: two types of Logistic Regression; Stochastic Gradient Descent; Random Forest; a linear Support Vector Machine; KNN; GaussianNB; and a Decision Tree. Using selected or default parameters in limited ranges did not in any of those methods appear to offer reliable classification.

### Example 2

For the second example, we also utilized a dataset from the North American Psychosis-Risk Longitudinal Study^13^. Here the goal is use of leukocytic microRNA (miRNA) levels to predict a subsequent conversion to frank psychosis in subjects meeting research criteria for clinical high-risk. Data included baseline levels of 130 leukocytic miRNA levels from 72 subjects, 37 who did not and 30 who did convert. The CALF solution uses the AUC as metric rather than the p-value used in the original publication^14^.

As shown in Figure 2 with the AUC metric and 100% of true data, CALF20 chooses only six predictors before ending; that is, no seventh predictor improves AUC; CALF6 has AUC 0.88. However, LASSO over a scan of s values can choose any number of weights up to 20. The CALF solution with empirical p-value 0.041 for the AUC metric is

CALF6 = +miR-941 -miR-25-3p -miR-378a-3p +miR-340-5p +miR-221-3p -miR-26b-3p

**Figure 2.**
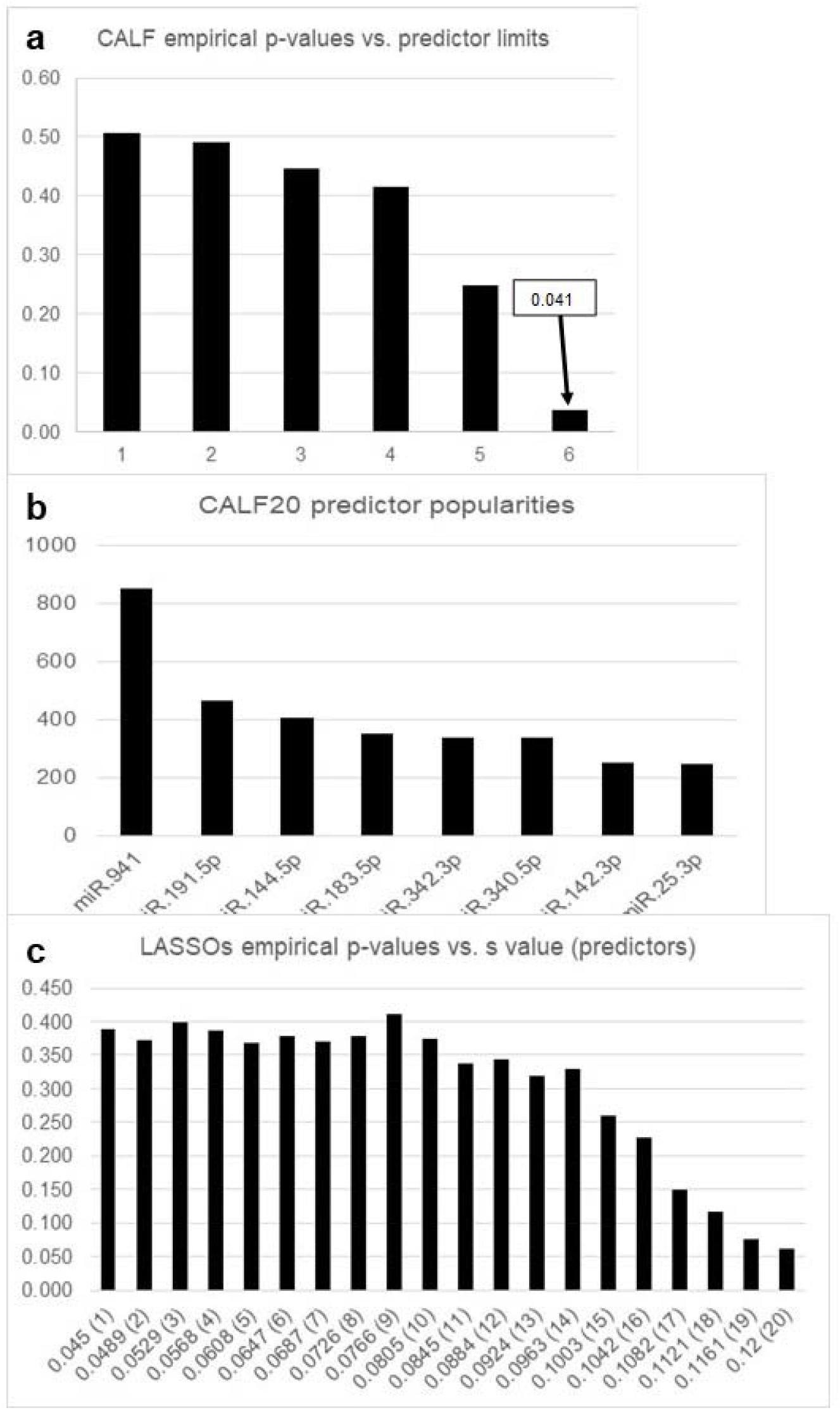
CALF *vs*. LASSO for example 2. (a) CALF is optimized at CALF6, where six is the limiting number of predictors used by CALF on 100% of true data. (b) There is a weak cliff beyond the first six predictors by popularity. (c) All LASSO solutions in the shown scan of s values have empirical p-value > 0.05.

### Comparison

The empirical p-values (0.041 *vs*. > 0.5) and the optimized number of predictors (6 *vs*. 20) for CALF6 and LASSO.12 imply CALF is a superior algorithm, given the constraint that at most 20 predictors as in the shown scan are used and the metric is AUC.

### Example 3

For the third example, we again utilized a dataset from the North American Psychosis-Risk Longitudinal Study^13^. Predictors are discrete but nonbinary, and there are more subjects than available predictors (N > p). Data included severity ratings of 19 symptoms from the Scale of Prodromal Symptoms (SOPS)^4,15^ determined at baseline from 72 subjects, 40 who did not and 32 who did develop a psychosis diagnosis. The process in this example differed from the original publication^8^ in that we utilized the same subjects as in examples 1 and 2, above, rather than the entire cohort.

As shown in Figure 3, symptom P1 (unusual thought content) was the most informative in the present analysis. Using only P1 yielded a poor empirical p-value of 0.050, thus illustrating the fact that goodness of fit in itself is not the best criterion for informativeness of a classifier. CALF4 and CALF9 had much better empirical p-values.

**Figure 3.**
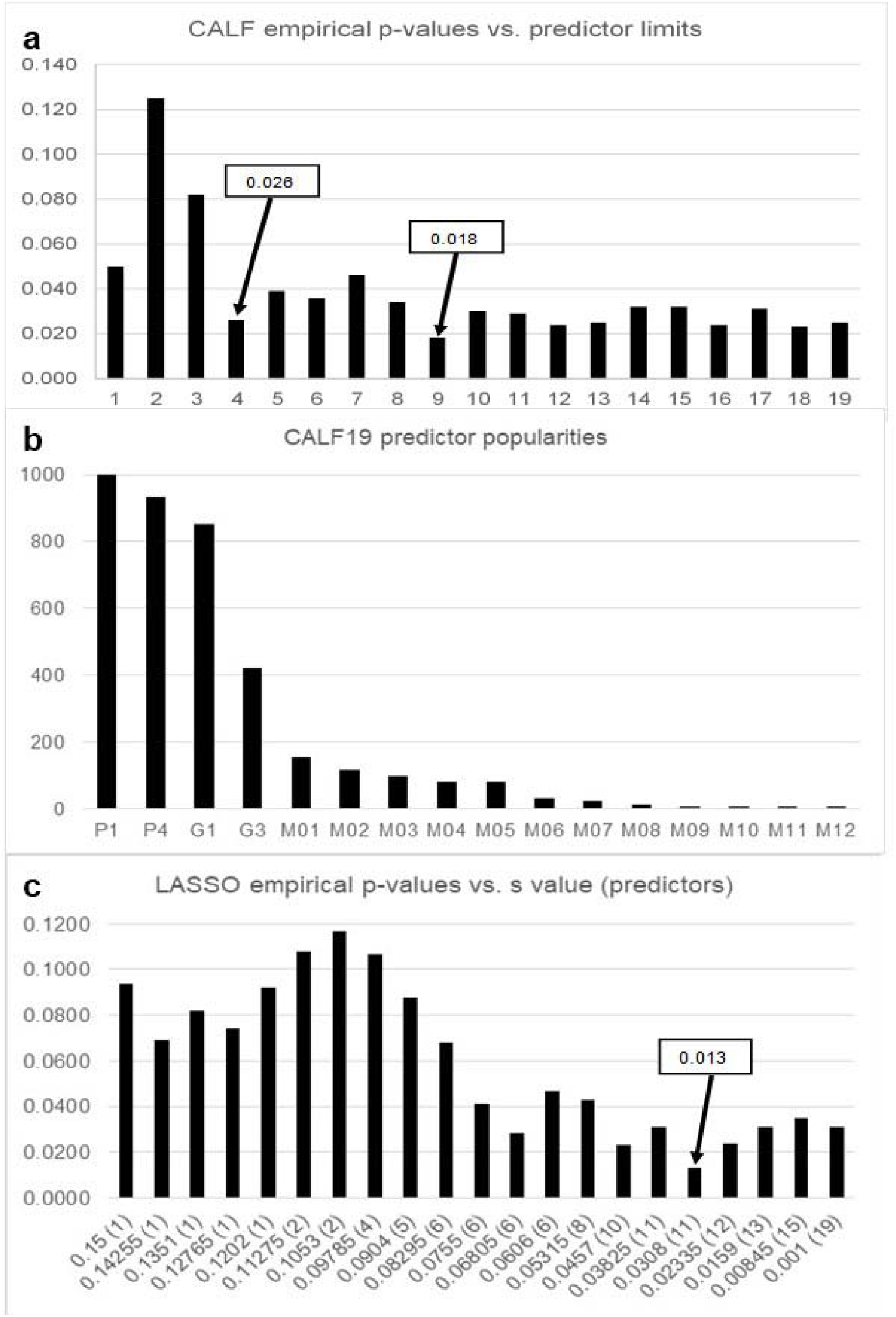
CALF *vs*. LASSO for example 3. (a) CALF4 empirical p-value (arrow) is 0.026 while CALF9 yields 0.018. However, the popularity profile (b) for CALF19 strongly suggests using L = 4. (Note three predictors were never chosen in 1000 random 90% subset experiments.) (c) Allowing s to scan from 0.15 (one predictor) down to 0.01 (19 predictors) shows LASSO.0308 with 11 nonzero weights (of 19 possible) has lower empirical p-value 0.013 but at the expense of more nonzero weights.

Hence, we select CALF and LASSO models

CALF4 = +P1 +G3 -P4 +G1

LASSO.0308 = +0.444 +0.147*P1 -0.125*P4 +0.105*G1 -0.063*N4 -0.052*P3 +0.050*G3 +0.048*D4 +0.027*N6 +0.012*P2 +0.007*P5 +0.006*G2

### Comparison

Note the four CALF predictors are among the 11 LASSO predictors with the same signs. The optimal LASSO solution LASSO.0308 has low empirical p-value (0.013) but uses 11 of the 19 predictors, suggesting overfitting. Valuing simplicity recommends selection of CALF4.

### Example 4

In this example 4 we utilized a dataset from Alzheimer disease (AD) research reported in the Biomarkers Consortium Project publication “Use of Targeted Multiplex Proteomic Strategies to Identify Plasma-Based Biomarkers in Alzheimer’s Disease” (https://fnih.org/our-programs/biomarkers-consortium/programs/plasma-based-biomarkers-alzheimers). The goal is use of blood plasma analyte levels to predict future development of dementia in persons meeting research criteria for mild cognitive impairment. Data included levels of 140 blood analytes from 378 subjects, 213 who were not and 165 who subsequently were diagnosed with dementia. The 140 blood analyte levels were determined from samples taken at the time of study enrollment. The graphs in Figure 4 recommend the CALF solution

CALF6 = APOC3 -CCL20 +Cortisol -NRCAM +MB -HBEGF

**Figure 4.**
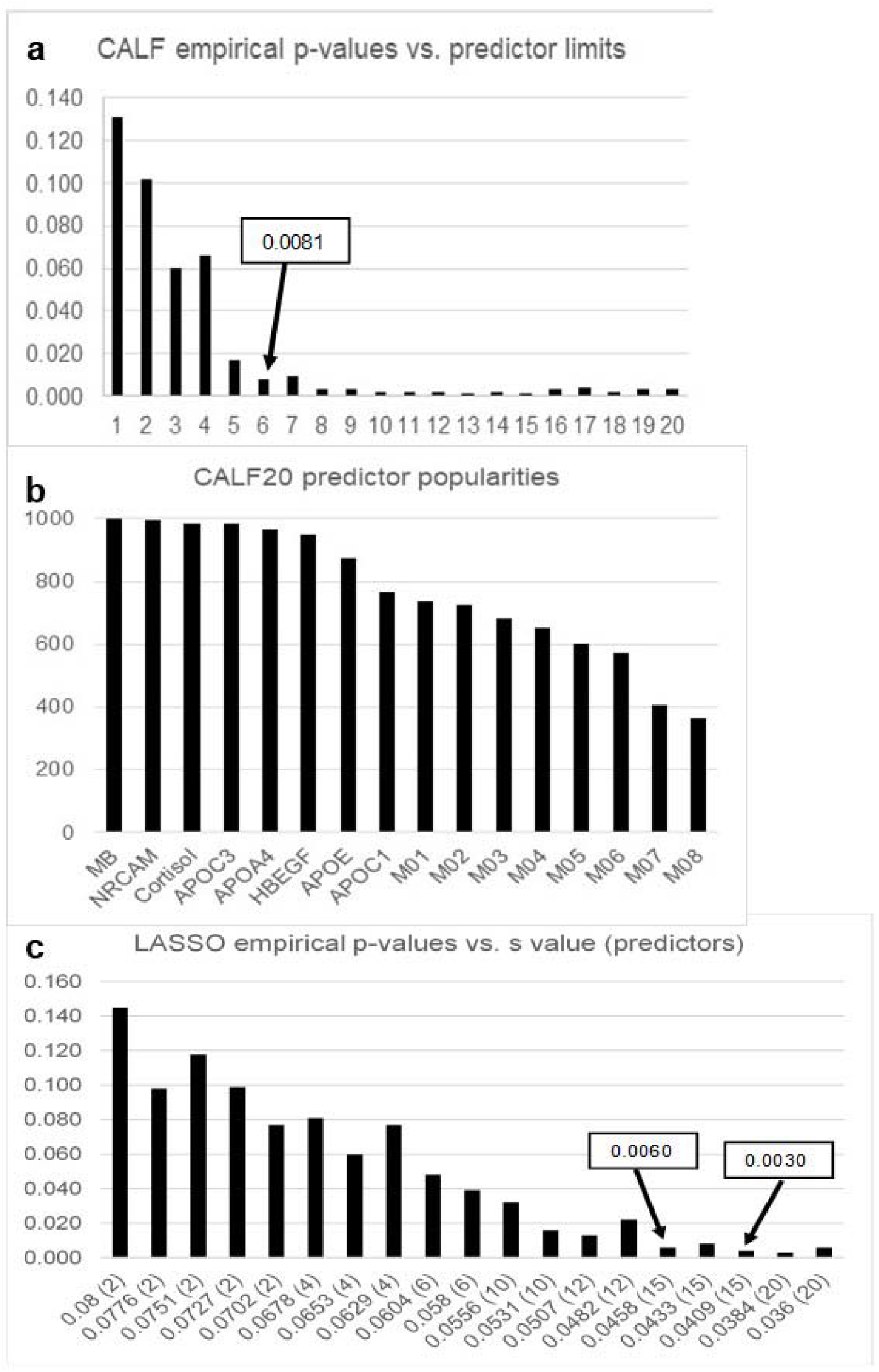
CALF *vs*. LASSO for example 4. (a) The first CALF low empirical p-value occurs with CALF6. (b) There is a weak cliff in popularity after the first six most popular predictors. Interestingly, among the eight most popular are four apolipoproteins (APOC3, APOA4, APOE, APOC1) (c) Low empirical p-values for LASSO require at least 15 predictors, risking overfitting.

### Comparison

Again, valuing simplicity recommends selection of the CALF solution.

### Example 5

Data are from the National Institute on Aging and Center from inherited Disease Research of Johns Hopkins University at https://www.ncbi.nlm.nih.gov/projects/gap/cgi-bin/study.cgi?study_id=phs000168.v2.p2. Data include age of onset (ranging from 52 to 98 years) from 582 AD patients. Predictors are minor allele/major allele coding at 9312 SNPs, scored as -1 (homozygous minor allele), 0 (heterozygous), 1 (homozygous major allele), followed by z-scoring. Age of onset itself is also recoded as a z-score. The CALF goal is selection of a ±1 weighted sum of a small set of SNPs that has a high Pearson correlation with age of onset. Although LASSO optimizes MSE, the correlation for any LASSO solution can be computed; likewise, the MSE for any CALF solution can be computed after OLS adjustment providing an intercept value and a common weight rescaling factor. These conversions allow direct comparisons of the two metric fits, correlation or MSE.

This data matrix is much larger, and so cross-validation analysis is feasible and thus included *infra*. Empirical p-value calculations for both CALF and LASSO applied to true data are very low. In fact, we have observed in 1000 random permutations of ages the true correlation from CALF with 1 to 20 predictors is ten or more standard deviations above averages of correlations of permuted data. Consequently, the results presented here do not included graphs of empiric p-values *vs*. number of predictors. Instead, correlations or MSEs are directly presented.

LASSO solutions jump from four to ten predictors for an s scan; no s values yield intermediate numbers of predictors. Going from two to three predictors and three to four yields improvements in MSE of -0.0082 and -0.0088 per predictor. However, going next from four (s = 0.2) to 10 (s = 0.19) predictors yields a smaller -0.0025 improvement per predictor. Altogether, these observations recommend CALF4 and LASSO.2, as shown in Figure 5. Also, LASSO.2 popularities include 16 SNPs with counts above 100 in 1000 trials, indicating relatively inconsistent or unstable LASSO.2 SNP choices relative to CALF4 in 90% subsets of true data.

**Figure 5.**
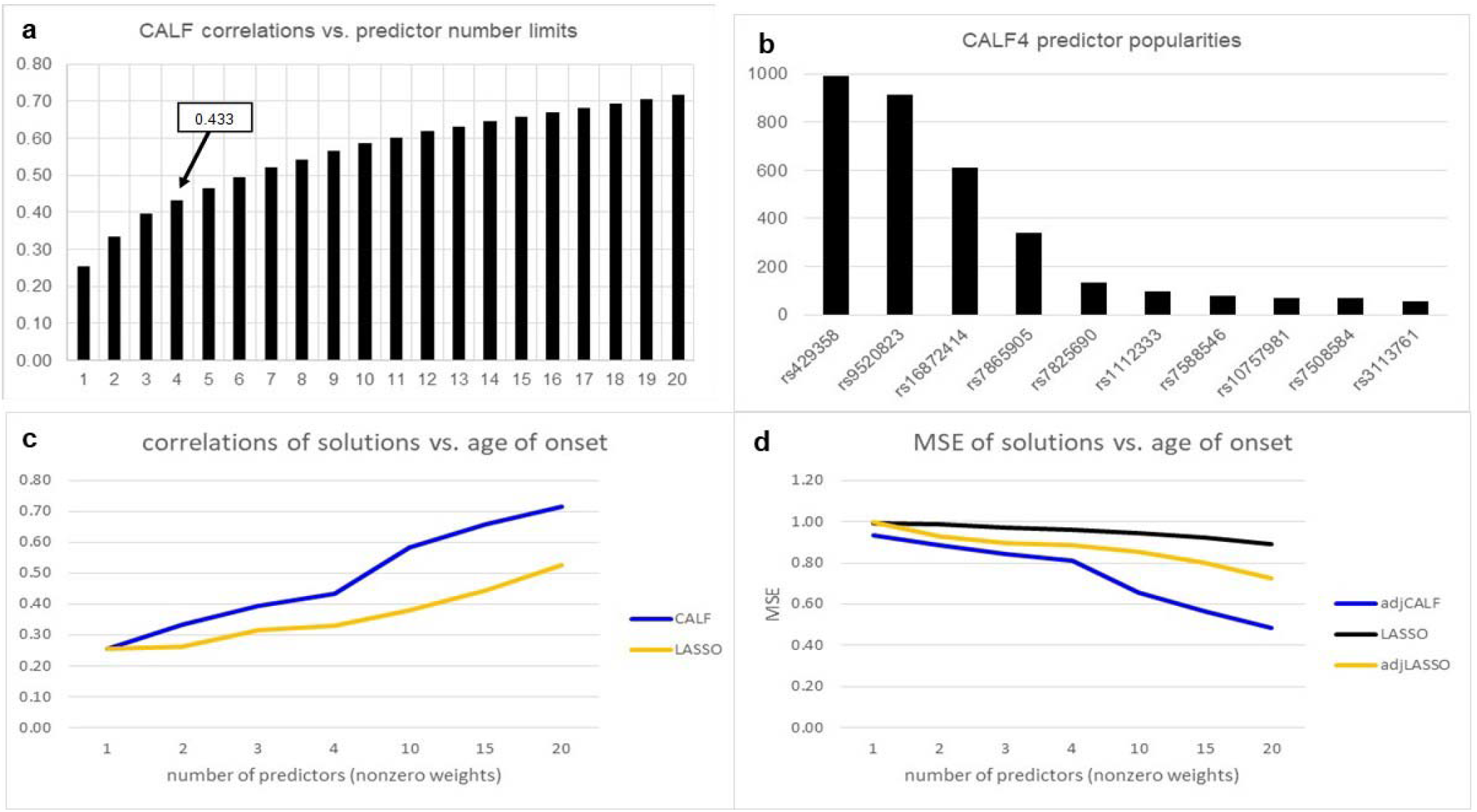
CALF *vs*. LASSO for example 5. (a) The correlations for CALFL over L from 1 to 20 increase from 0.254 to 0.717 with 0.433 for CALF4. (b) A cliff in CALF popularities after four predictors recommends L = 4. In 1000 90% subset trials there were only five SNPs with counts above 100. (c) Regarding correlations, CALF consistently outperformed LASSO (no s values allowed certain numbers of nonzero weights). (d) Regarding MSE over 1 to 20 predictors, adjusted CALF outperformed both basic LASSO and adjusted LASSO. For LASSO.2 with four predictors there were 16 SNPs with popularity counts of at least 100. The difference *vs*. CALF SNP popularities suggests the CALF choices are more consistent and stable.

The CALF4 solution (with predictors in order of CALF selection) is

CALF4 = -rs429358d +rs9520823 +rs16872414 -rs7865905

Interestingly, we found CALF4 applied 1000 times to data with randomly permuted age of onset yielded average correlation = 0.299 with standard deviation (sd) 0.013 in a normal distribution. Thus, the true CALF4 correlation 0.433 is about 10 sds greater than the average CALF4 correlation.

Individually, the correlations of the first two SNPs are -0.25 and +0.23, the highest in magnitudes of all 9312 SNPs. As expected, the leading SNP rs429358 (https://www.ncbi.nlm.nih.gov/snp/?term=rs429358) is one of two SNPs that define the APOE _ε_4 variant, a well-known genetic risk factor for AD. The protein APOE is a lipid carrier in the central and peripheral nervous systems^15^. The APOE _ε_4 variant is said to have “unparalleled influence on increased late-onset AD risk”^16^ (late-onset predominates in AD). The SNP rs429358 has alleles T>C and is in exon 4 of gene APOE https://www.ncbi.nlm.nih.gov/snp/rs429358. In our normalization system (0/0 ← -1, 0/1 ← 0, 1/1 ← +1, thence to z-scores), the C/C allele maps to a negative z-score. Thus, the CALF4 weight -1 is consistent with increasing CALF4 function values with age and hence, stronger correlation with age of onset and association with risk of late-onset AD, as expected.

For comparison, the LASSO.2 solution with four weights is

LASSO.2 = -3.61E-06 -5.23E-02*rs429358 +2.67E-02*rs9520823 -1.67E-03*rs7865905-9.91E-04*rs8125417

The four weights are ordered by decreasing magnitude. Note that first three LASSO.2 predictors are also among the four predictors in CALF4 in the same order and with the same signs. However, only six predictors are in both the 20 predictors of CALF20 and the 20 predictors of LASSO.17, underscoring the differences in the algorithms’ predictor choices. In particular, LASSO.2 simply chooses the four predictors with strongest individual correlation magnitude.

Continuing with the constraint of using four predictors, a direct comparison over true data yields superior CALF results: correlation CALF4 = 0.433; LASSO.2 = 0.332. Likewise, after OLS adjustment of both solutions, the MSEs are: adjCALF4 = 0.811; adjLASSO.2 = 0.821. Thus, after adjustments, CALF4 outperforms LASSO.2 in both metrics.

As a final note on this example, the second-most popular SNP in both CALF4 and LASSO.2 solutions was rs9520823; the 1/1 variant was associated with later age of onset (yielding a positive regression weight). This SNP is in an intron of gene ABHD13, a ubiquitously expressed gene in the ABHD family with protein products important in lipid synthesis and degradation^17^. If the rs429358 SNP is deleted from the data matrix, rs9520823 becomes first-chosen and CALF4 correlation decreases from 0.433 to 0.421 (data not shown). That is, after discarding APOE, CALF4 leads with the SNP for ABHD13 and correlates almost as well with age of onset. Investigation of causality involving APOE and ABHD13 functions in AD is suggested by the above findings. Lipid transport, reception, and metabolism are active AD research arenas^18^.

### Example 5

To illustrate cross-validation performance we applied CALF with a limit of four nonzero weights to 1000 random 90% subsets of true data for training and then applied each of those 1000 solutions to the 1000 complementary 10% subsets, recording the correlation values. The distribution of 1000 cross-validation results is shown in Figure 6.

**Figure 6.**
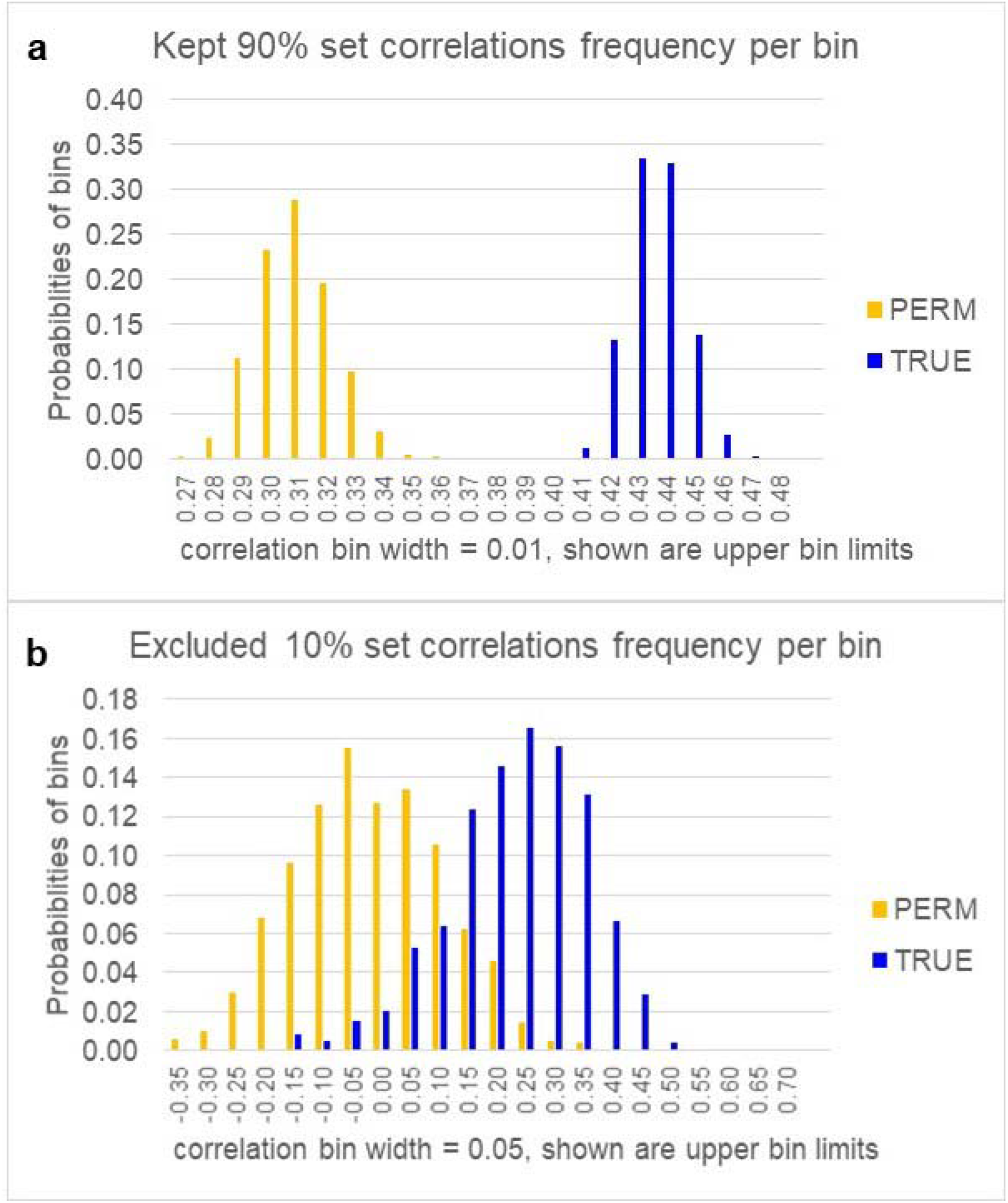
Cross-validation analysis. (a) A histogram for cross-validation CALF4 on random 90% kept subsets of SNP data in 1000 trials includes bins of correlation values *vs*. ages of onset. TRUE bars denote values of correlations using 1000 random (with replacement) 90% subsets. PERM bars denote first a random permutation of all 582 ages of onset, then a repetition of the TRUE analysis. Thus, PERM is the null hypothesis result, that is, the absence of information. Welch t-test p-value of the two sets of correlations is ∼0.00. (b) Shown is a histogram of bins of correlation values with ages of onset. TRUE bars denote values of correlations using 1000 random 10% subsets complementary to each of the subsets in (a). PERM denotes first arandom permutation of all 582 ages of onset, then a repetition of the TRUE calculation. Thus, PERM is the null hypothesis result. Normal distributions of these TRUE and PERM excluded values have K-S goodness of fits α = .0.05 and 0.20 (so the PERM distribution is very close to normal). The average of PERM correlations in the trials was ∼0.006, but the average of TRUE correlations was ∼0.26. Assuming normality of both sets, Welch t-test p-value (two-sided) is ∼5.45E-307.

### Comparison

We conclude that CALF4 achieves cross-validation in the permutation test sense for this example. Because using the s parameter in LASSO does not generally determine the number of nonzero weights in random subset solutions, analogous analysis of LASSO demanding simplicity (a fixed number of nonzero weights) is not possible.

## Discussion

Components of the CALF function are geometrically suggested by Figure 7a.

**Figure 7.**
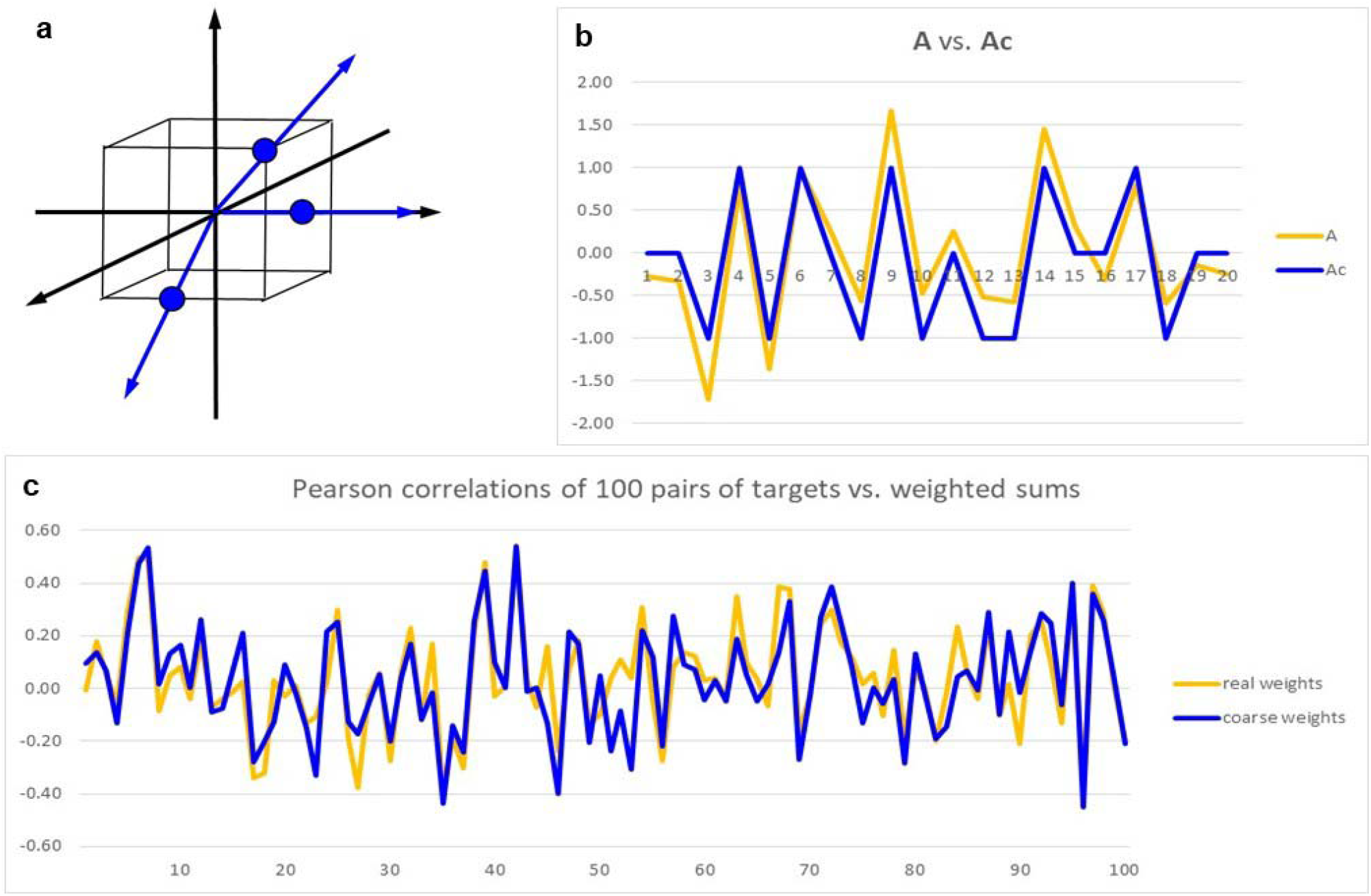
Geometry of CALF and simulations of coarse approximations. (a) Three rays (of 26 possible) employed by CALF in 3-dimensional space relate to coordinates of the vertices of the cube (the eight triplets of ±1). One ray passes through a vertex, one through the midpoint of an edge, and one through the midpoint of a face. (b) Visual representation of a typical, single coarse approximations of a real 20-dimensional vector. The real vector **A** has 20 normally distributed components, mean = 0 and sd = 1; components of the coarse approximations **Ac** use threshold = ±0.4307. That is, if a component of **A** is < 0.4307, then the corresponding component of **Ac** is -1, and so on. (c) Among 20-dimensional vectors, a plot of 100 comparisons of dot products of **A** with **B** vs. **Ac** with **B**, where **A** and **B** are normally distributed and **Ac** is the coarse approximation of **A**.

The surface area of the cube in Figure 7a is 24, so the ratio of the number of possible CALF rays to area is 26/24. Generalizing the concepts to n-dimensional space, there are 3^n^-1 rays passing from the origin through all points with coordinates that are combinations ±1 or 0 (excepting the origin itself). The (n-1)-dimensional area of the surface of the n-cube is n2^n^. Thus, the ratio of the number of all possibly CALF rays to surface area is (3^n^-1)/(n2^n^). It can be shown using calculus that in dimensions > 4, this ratio exceeds 1.05^n^. Roughly speaking, the rays available to CALF approximations, each representing a direction available for the approximation, become “exponentially crowded” on the surface of the n-cube as dimension increases.

Figure 7b shows that normally distributed vector components are fairly tracked by coarse approximations, implying the directions of the two vectors in 20-dimensional space are similar. Figure 7c shows that correlations of a normally distributed vector or its coarse approximation with an independent 20-dimensional vector (such as a dependent vector in a regression analysis) are little different.

In any linear regression model, an approximation of the i^th^ entry in the n-dimensional dependent vector is the dot product of the weight vector and the j^th^ vector of p predictor vectors. If components of the weight vector are selected from ±1 or 0, then in, say, 20-dimensional space, 3^20^-1 = ∼3.49E9 selections are possible. A coarse weight vector can be thought of as an approximation of an exact, optimal, real-valued *direction* (ray) in 20-dimensional weight space. For the Welch p-value, AUC, or correlation scores, only the *direction* of the weight vector, not its magnitude, is significant; positive magnitudes are irrelevant for those metrics.

## 5 Conclusions

CALF deterministically seeks a coarse sum of a few predictors to optimize a score. As with any multiple regression approach, the goal of CALF is discovery of a network of informative predictors, not identification of single marker as individually informative. We can do so despite the computational explosion of numbers of possible sets of subsets precisely because CALF uses a greedy approach. Successful permutation tests and other tests then provide model quality assessment.

The examples show that CALF sometimes finds a simpler regression with superior metric performance *vs*. basic LASSO. Thus, there might be cases in which CALF finds statistically significant information in data while LASSO finds only a trend. As noted above, it appears that using LASSO without considering CALF risks wasting resources.

## Methods and Implementation

### R Version

An implementation of the CALF algorithm in the R language is available through the Comprehensive R Archive Network (CRAN) as package `CALF` implemented as the function `calf()`. The `calf()`function may be run with a binary or nonbinary dependent vector (`targetVector`). In the binary case, `calf()` seeks to optimize Welch t-statistic p-value or AUC (`optimize = pval` or `optimize = AUC`). Due to the symmetries of those optimizations, the algorithm chooses the initial weight to be +1. According to user preference of control = 0, case = 1 or the opposite, the weights in the final sum may be reversed. For subsequent MSE optimization, a simple linear transformation may be applied (which of course does not alter p-value or AUC performance).

Alternatively, for a dependent vector that is real-valued, nonbinary `calf()` is employed for a positive Pearson correlation using `optimize = corr` (correlation). Again, a linear transformation may subsequently be applied to minimize MSE (but preserve correlation). The initial weight might be +1 or -1. The CALF User Guide fully documents this binary *vs*. nonbinary difference as well as other aspects of the `calf()` function.

Four supplementary functions are also provided. Permutation tests randomly permute entries in the dependent vector to reveal the empirical p-value. CALF may be applied to many random subsets (of one or other fixed fraction of all subjects) to find the most “popular” predictors, displaying tables of choices and performance values. Another function `cv.calf()` enables cross-validation, repeated and/or stratified. For binary dependent data it selects random subsets of control data of fixed proportion and random subsets of case data of the same proportion; for continuous dependent data, it selects a fixed proportion of all data. Then cross-validation computes CALF weights and applies the resulting weighted sum to the complementary set. Documentation for these supplementary functions is included in the CALF User Guide in the package. The function `perm_target_cv()` will conduct the same procedure as cross-validation, documented above, however it will permute the target column (dependent vector) of the data as a very first step, usually the first column, with each iteration of the process.

Presently, our scope for `CALF` was simply to provide an accurate implementation of the CALF algorithm plus common methods of evaluation of regression sums. There certainly is room for improvements and enhancements, but changes will include support of the current functional interfaces; thus, there should be no function deprecation with future versions. Further, source code may be minimized such that existing redundant functionality will be moved to a single function, thus making code more concise and maintenance more streamlined. An important future goal is a version that is conformant to popular R statistical modeling packages, especially `caret`.

### Python Version

A Python implementation of CALF obtainable via the PyPi repository at https://pypi.org/project/calfpy/1.16.0/. In a Python environment, installation follows typing `pip install calfpy` at the command line. The Python version functions in the same manner as the R version, as described above. Given that Python does not natively offer a data frame structure or mechanisms to operate on data frames, the Python CALF implementation relies upon the pandas, `numpy`, and `scipy` packages to handle such.

Some hallmark programming techniques often employed in Python are only minimally used in this implementation, e.g., list comprehension. The purpose for this break from style was two-fold: to ensure non-Python programmers could more easily review the code, if desired; and to ensure the code remains somewhat in step with the existing R version. As the core processing is mainly done by the packages listed above, it is not believed these style changes affect performance significantly.

### Data

Sample data that may be used for duplication of all the stated results is available via GitHub as an unrestricted supporting resource at https://github.com/jorufo/CALF_SupportingResources.

## Acknowledgement

Ms. Stephanie Lane used Excel spreadsheets with macros from CDJ to create the first R code for CALF; she is acknowledged and listed as creator in CRAN.org documents.

