## Supplement 1 for "Frisky CALF sometimes outruns LASSO"

Supplement: Suboptimal CALF solutions

The CALF algorithm is greedy and as such looks forward only one step. That is, in seeking a metric improvement, CALF considers adding only one more marker per algorithmic cycle. An intermediate or terminal sum obtained by such steps can be called “reachable” by CALF. A sum with a metric value at least as good as all others (of 3^n – 1 possible, discarding the all 0 sum) can be called “optimal.” (Of course, pairs of sums such as +M1+M2+...+Mn and -M1-M2-...-Mn have the same metric values.) In the following examples, the metric is taken to be the Student t-test p-value (p-val). CALF initially biases the very first coefficient to be +1; if the final CALF sum is a function with undesirable sign of correlation to the target data, then signs of all the nonzero coefficients of the sum can be reversed.

It is not true that CALF always finds the optimal sum, that is, the best sum of marker values with ±1 coefficients for approximation of trends in target data, the optimal sum, can be unreachable.

Table1. A toy data matrix for which binary CALF with p-value metric yields a suboptimal solution.

| case/ctrl | M1 | M2 | M3 |
| --- | --- | --- | --- |
| 0 | 0.224 | 0.494 | 0.256 |
| 0 | 0.096 | 0.433 | 0.131 |
| 0 | 0.607 | 0.010 | 0.783 |
| 1 | 0.444 | 0.499 | 0.082 |
| 1 | 0.834 | 0.226 | 0.935 |
| 1 | 0.430 | 0.821 | 0.136 |
| 1 | 0.163 | 0.219 | 0.822 |

CALF first chooses M1 as the single best marker, then, of ±M2 or ±M3, CALF selects +M2 as the best marker to add, forming +M1+M2. Attempting to add ±M3 worsens the metric value, so CALF terminates with +M1+M2 (p-val = 0.22). However, the sum 0*M1+M2+M3 = M2+M3 has a better metric value (0.17); the optimal sum +M2+M3 is unreachable by CALF.

Table2. A second toy data matrix for which binary CALF with p-value metric yields a suboptimal solution.

| case/ctrl | M1 | M2 | M3 | M4 |
| --- | --- | --- | --- | --- |
| 0 | 0.738 | 0.954 | 0.770 | 0.200 |
| 0 | 0.885 | 0.457 | 0.650 | 0.963 |
| 0 | 0.569 | 0.943 | 0.960 | 0.613 |
| 1 | 0.522 | 0.728 | 0.700 | 0.755 |
| 1 | 0.277 | 0.451 | 0.000 | 0.623 |
| 1 | 0.214 | 0.759 | 0.200 | 0.300 |
| 1 | 0.285 | 0.751 | 0.235 | 0.931 |

Again, CALF first chooses M1 as the single best marker, then, from ±M2, ±M3, or ±M4, CALF selects +M2 as the best marker to add, forming +M1+M2 as a classifier function (p-val = 0.018). Attempting next to add ±M3 or ±M4 worsens the metric value, so CALF terminates with +M1+M2. However, the unreachable sum +M1-M2+M3-M4 is superior (p-val = 0.012).
