## Supplement 2 for "Frisky CALF sometimes outruns LASSO"

Supplement: Application of selected conventional models to binary classification problem

The purpose of this supplement is to compare CALF performance for the binary example with performances of several other algorithms. For the other algorithms, default values or a limited set of tuned parameters were used.

That is, in addition to CALF and LASSO as described in the text, this supplement displays results of several classification schemes that were applied to the same example (135 normalized blood analytes as measured in 40 controls and 32 cases). The csv is in Supplement 1. The results are representative of the additional types of models but of course are not exhaustive descriptions of results of all possible tuned parameters or classifier models.

To further explore conventional classifier models, we applied nine different types of algorithms.

For each of the models developed below, zero, one or two model parameters were optimized using a simple 10000 cross validation (CV) procedure. Specifically, 20% validation hold out sets (8 of 40 controls and 6 of 32 cases) were generated 10000 times. Depending on the model, either a dense grid-search strategy or a random (statistical) search was performed. For each of the chosen models, the estimated parameters and the mean accuracy metric for the CV using the best parameters estimates are reported. Default values were used for all other parameters. The top five important predictors for the model are also reported since, by comparison, CALF selected five of 135 predictors with optimized surpass rate. Depending on the model, the predictors may be actual coefficients or estimates of importance.

To test the significance of the fit beyond the 10000 CV experiments, a permutation test was performed using the (crudely) optimized estimated parameters. The permutation procedure was executed 10000 times. The associated p-values (surpass rates)^1^ are reported in the following tables. These provide an estimate of the likelihood that the performance of the discovered classifier is due merely to chance. Also, the value of the AUC score is reported for each model.

Models: Basic usage

All the following models were accessed through the Python package Scikit-learn^2^.

1) A Logistic Regression Classification model using an elastic net penalty function was tested.^3^

The elastic net penalty parameter I1_ratio was optimized using a discrete grid search in the range (0, 1).

2) A second Logistic Regression Classifier using a pure Lasso penalty^4^ was also performed. Default parameters.

3) A Stochastic Gradient Descent (SGD) method^5^ was applied. An elastic net penalty was applied and the l1_ratio was grid searched in the range (0,1).

4) A Random Forest Classifier (RFN) model^6^ was applied. The only parameter optimized using a grid searching scheme was the number of estimators.

5) A linear Support Vector Machine (SVM) model^2^ was applied. A randomize search was used to estimate the parameters gamma (reciprocal sampling: 0.001 to 0.1) and the scale factor C (uniform sampling 1 to 10).

6) A KNN model^7^ was applied with grid search optimization of the leaf size (5,40) and the number of neighbors (10,70).

7) GaussianNB as in Scikit-learn^2^ using all default parameters was applied.

8) Decision Tree^8^ using all default parameters was applied.

9) Lastly, a hard voting consensus^9^ as available from Scikit-learn was constructed from all the models.

None of the models using all 135 predictors showed significant fit quality based on the CV scores and the AUC of ROC curves. A simple consensus method (hard voting) of all the tests also failed. A summary appears in Table 1.

Table 1. Selected results for a suite of models applied to the binary classifier problem.

| Model (parameters) | Score (%) | Perm p-value | AUC |
| --- | --- | --- | --- |
| SGD (alpha=0.6) | 58.75 | 0.04 | 51.48 |
| Logistic Regression (elastic net) (alpha=0.0) | 58.57 | 0.13 | 55.70 |
| SVC linear(C=4.75, gamma=0.71) | 54.29 | 0.29 | 58.20 |
| Logistic Regression (purely lasso) | 50.00 | 0.52 | 54.29 |
| Random Forest (estimators=10) | 51.79 | 0.86 | 45.54 |
| KNN (leaf size=10, neighbors=10) | 62.86 | 0.02 | 51.56 |
| Gaussian NB | 41.79 | 0.88 | 40.93 |
| Decision Tree | 51.61 | 0.54 | 45.62 |
| Hard voting consensus | 53.04 | NA | NA |

PCA and ICA predictor set reduction

To examine the importance of reducing the dataset dimension prior to application of the models, both Principle Component Analysis^10^ (PCA) and Independent Component Analysis^11^ (ICA) preprocessing were performed. For ICA, the predictors were whitened. The PCA and ICA methods were accessed from the Scikit-learn package. For both approaches, a total of 5 components were requested. From these, a set of models were applied including the parameter optimization step, followed by permutation testing and the AUC-ROC determination. The results are collected in the following Tables. Generally, reduction of the components had little effect on the final results. In all cases poor classifier performance continued to be observed. In the following plot are the PCA eigenvalues. Clearly, reduction of predictors based on either variance (PCA) or information (ICA, up to 500 iterations) is insufficient to build a quality model for this small data set.

Table 2. Principal component analysis (PCA) with five components specified.

| Model (parameters) | Score (%) | Perm p-value | AUC |
| --- | --- | --- | --- |
| SGD (alpha=0.9) | 55.71 | 0.034 |  |
| Logistic Regression (elastic net, alpha=0.9) | 57.14 | 0.251 | 0.593 |
| Logistic Regression (LASSO) | 55.89 | 0.315 | 0.489 |
| Random Forest (estimators = 24) | 50.00 | 0.739 | 0.496 |
| Decision Tree | 41.61 | 0.933 | 0.402 |
| Hard voting consensus | 57.32 | NA | 0.403 |

The PCA eigenvalue spectrum indicates that the amount of variance explained when including only 5 components is rather small, ranging from 0.105 down to 0.048.

In the next Table 3 are analogous tests using Independent Component Analysis (ICA) for predictor set reduction.

Table 3. Independent component analysis (ICA) using five components

| Model (parameters) | Score (%) | Perm p-value | AUC |
| --- | --- | --- | --- |
| SGD (alpha=0.2) | 50.00 | 0.034 | 0.465 |
| Logistic Regression (elastic net, alpha=0.1) | 55.71 | 0.251 | 0.372 |
| Logistic Regression (LASSO) | 55.71 | 0.315 | 0.491 |
| Random Forest (estimators = 24) | 50.00 | 0.739 | 0.430 |
| Decision Tree | 38.57 | 0.933 | 0.534 |
| Hard voting consensus | 52.86 | NA | NA |

Limited predictor set computation

A (biased) test of the five best predictors as selected by the CALF solution on 100% of true data (+M040+M135-M123+M070-M086) was also performed using several of the tested traditional models. In all cases substantially better CV scores, permutation p-values, and AUC-ROC were observed. This may suggest that the predictors identified by CALF do indeed capture relevant association of predictors and the target.

Table 4. Using the five (best) CALF predictors only

| Model (parameters) | Score (%) | Perm p-value | AUC |
| --- | --- | --- | --- |
| SGD (alpha=0.2) | 67.86 | 0.0001 | 0.755 |
| Logistic Regression (elastic net, alpha=0.0) | 76.43 | 0.0002 | 0.834 |
| Logistic Regression (LASSO) | 76.43 | 0.0001 | 0.827 |
| Random Forest (estimators = 19) | 66.43 | 0.0009 | 0.695 |
| Decision Tree | 57.14 | 0.364 | 0.603 |
| Hard voting consensus | 76.43 | NA | NA |

A representative ROC plot for the Logistic Regression model using the five predictors is in Figure 1.

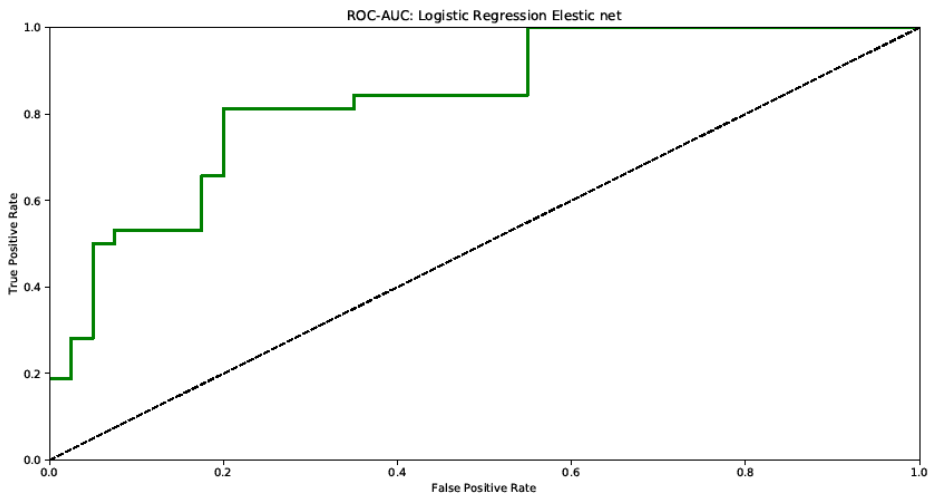

Figure 1. ROC of a Logistic Regression model built with the five predictors selected by CALF.
